# Bacterial Glycogen as a Durable Energy Reserve Contributing to Persistence: An Updated Bibliography and Mathematical Model

**DOI:** 10.1101/536110

**Authors:** Liang Wang, Qinghua Liu, Xinle Tan, Ting Yang, Daoquan Tang, Wei Wang, Michael J. Wise

## Abstract

Glycogen is conventionally viewed as a transient energy reserve that can be rapidly synthesized for glucose accumulation or mobilized for ATP production and blood glucose homeostasis in higher organisms. However, this understanding is not completely applicable to prokaryotes due to glycogen structural heterogeneity. A number of studies have noted that glycogen with short average chain length *g_c_* in bacteria has the potential to degrade slowly, which might prolong bacterial survival in the environment and thus enhance potential for transmission to new hosts. This phenomenon has been examined over the past few years and called the durable energy storage mechanism hypothesis (DESM). In this updated bibliography, we summarize recent progress and provide a mathematical model of glycogen as a durable energy reserve.

## Introduction

Glycogen is a highly branched, water-soluble, and homogeneous polysaccharide that is widely presented in all three life domains. It consists of α-1,4- and α-1,6-linked glucosyl units only. Conventionally, glycogen is linked with maximized storage of glucosyl residues and quickly mobilized glucoses for energy supply and blood sugar homeostasis in cellular metabolism of higher organisms (1). However, recent studies indicate that some bacterial glycogen has the potential to function as a durable energy reserve (2–5). This feature is largely regulated by average chain length *g_c_*, one of the determinants of glycogen structure. A set of experimental measurements through the periodate oxidation method has confirmed that *g_c_* in muscle and liver glycogen is comparatively consistent, ranging from 11 to 15 d.p. (degree of polymerization) across eukaryotic species (2). In contrast, *g_c_* in bacteria is highly divergent within a range of 6.6-21 d.p. (2). Thus, glycogen structure between eukaryotes and prokaryotes could be different and may have impacts on their functions. Correlation analysis has already shown that glycogen with shorter *g_c_* acts as a slow-degrading energy reserve and contributes to prolonged bacterial survival in hash environments. For example, enzymology analysis finds that shorter *g_c_* means higher percentage of α-1, 6- branching points, which requires more energy to break down (2). In addition, branching points are thermodynamically favored during formation. Thus, highly branched glycogen is more structurally stable (3). Moreover, acid hydrolysis confirms that α-1, 4- glycosidic linkages break up at a faster rate than α-1, 6- glycosidc linkages (4, 5). Thus, glycogen durability could be an inherent characteristic, rather than relying on enzyme-substrate interactions (5). In sum, slow-degrading glycogen provides a passive energy-saving strategy for bacteria to adapt to hash conditions, such as nutrient deprivation. A hypothesis of glycogen as a durable energy reserve has been termed as Durable Energy Storage Mechanism (DESM) to explain the long-term survival of some bacteria in harsh environments (2).

Since the proposal of glycogen as a representative of the DESM hypothesis in 2011, a variety of studies have focused on the mechanism (Table 1). However, so far, there is no experimental confirmation of the existence of glycogen as a durable energy reserve. (6) attempted to use in situ progressive truncation of GBE N-terminus to alter glycogen *g_c_* in order to investigate how *g_c_* can influence bacterial environmental persistence. This experiment confirmed the feasibility of *g_c_* manipulation through gene truncation, and the corresponding impacts on Escherichia coli stress resistance and biofilm formation were observed. However, there was a fundamental flaw of the experiment: the linkage of glycogen amount and *g_c_*, which left the conclusion of the study difficult to illustrate. In addition, the foundation of glycogen as DESM is also compromised if we consider that the variability of *g_c_* in bacteria might be caused by different methods performed by independent groups (Table 2). In this update, we assess recent progress toward understanding glycogen as a durable energy reserve in bacteria. We then investigate the classical glycogen model initially proposed by Whelan from a novel perspective in order to illustrate how *g_c_* can influence glycogen structure theoretically. Finally, the relationship between glycogen metabolism and bacterial durability is comprehensively explored in order to provide further support for the hypothesis.

**Table 1:** Studies citing the Durable Energy Storage Mechanism (DESM) hypothesis since its first proposal (2)

| No | Year | Species | Argument | Ref |
| --- | --- | --- | --- | --- |
| 1 | 2012 | *Synechocystis* sp. PCC 6803 | Short gc increases durability | (7) |
| 2 | 2012 | *Agrobacterium tumefaciens* | Short gc increases durability | (8) |
| 3 | 2013 | *Rhodococcus* sp. | Glycogen for carbon starvation | (40) |
| 4 | 2013 | *Lactobacillus acidophilus* | Glycogen for diverse lifestyle | (41) |
| 5 | 2013 | *Corynebacterium glutamicum* | Exponential utilization of glycogen | (42) |
| 6 | 2013 | *Escherichia coli* | Glycogen supports viability | (43) |
| 7 | 2013 | *Escherichia coli* | Glycogen supports viability | (44) |
| 8 | 2013 | *Escherichia coli* | Short gc increases durability | (45) |
| 9 | 2014 | General bacteria | Intracellular energy storage | (46) |
| 10 | 2014 | *Lactobacillus acidophilus* | Glycogen for diverse lifestyle | (47) |
| 11 | 2014 | Fecal microbiota | Further study needed | (9) |
| 12 | 2015 | *Corynebacterium glutamicum* | Glycogen accumulation | (48) |
| 13 | 2015 | *Lactobacillus* | Glycogen for diverse lifestyle | (16) |
| 14 | 2015 | *Bifidobacterium* | Glycogen for diverse lifestyle | (15) |
| 15 | 2015 | *Escherichia coli* | GBE for optimal structure | (49) |
| 16 | 2015 | *Escherichia coli* | Glycogen supports viability | (50) |
| 17 | 2015 | *Escherichia coli* | Short gc increases durability | (6) |
| 18 | 2015 | *Vibrio vulnificus* | Short gc increases durability | (10) |
| 19 | 2016 | *Galdieria sulphuraria* | Controversial | (12) |
| 20 | 2017 | *Methyloprofundus sedimenti* | Glycogen as an energy storage | (51) |
| 21 | 2017 | *Galdieria sulphuraria* | Slow degradation of short gc glycogen | (13) |
| 22 | 2017 | General bacteria | Slow degradation of short gc glycogen | (14) |
| 23 | 2017 | *Escherichia coli* | Glycogen supports viability | (52) |
| 24 | 2017 | General bacteria | Durable energy storage mechanism | (53) |
| 25 | 2018 | General bacteria | Polyphosphate metabolism | (54) |
| 26 | 2018 | *Vibrio vulnificus* | Short gc increases durability | (11) |

**Table 2:** Methods used for measuring glycogen *g_c_* in bacteria and eukaryotes (adapted from (2) with newly sourced methods)

| Bacteria | $g_c$ | Methods | Ref |
| --- | --- | --- | --- |
| *Aerobacter aerogenes* | 13 | β-amylolysis, Methylation | (55, 56) |
| *Agrobacterium tumefaciens* | 13 | N/A | (2) |
| *Arthrobacter globiformis* | 6.6 | β-amylolysis, Methylation | (57) |
| *Arthrobacter* spp | 7-9 | β-amylolysis | (57, 58) |
| *Bacillus megaterium* | 10-11 | N/A | (2) |
| *Bacillus stearothermophilus* | 21 | Periodate oxidation | (59) |
| *Chromatium* strain D | 11.1-11.4 | Periodate oxidation | (60) |
| *Clostridium botulinum* | 17 | Periodate oxidation | (61) |
| *Desulfurococcus* | 7 | Periodate oxidation | (62) |
| *Escherichia coli* | 12,10.8,13,14 | Isoamylase debranching and chromatography (10.8), High-Performance Anion-Exchange Chromatography | (63, 64) |
| *Klebsiella pneumoniae* | 11.6 | Isoamylase debranching and chromatography | (63) |
| *Mycobacterium tuberculosis* | 7-9,10-12 | N/A (7-9), Oxidation (10-12) | (2, 65) |
| *Neisseria perflava* | 11-12 | N/A | (2, 66) |
| *Nostoc muscorum* | 13 | Periodate oxidation | (58, 67) |
| *Prevotella ruminicola* | 8 | Periodate oxidation | (68) |
| *Pseudomonas* V-19 | 8 | Methylation, Periodate oxidation | (69) |
| *Selenomonas ruminantium* | 12, 23.5 | Liquid chromatography (12), Methylation (23.5) | (70, 71) |
| *Sphaerotilus natans* | 11.5, 13 | Inferred from H1-NMR (11.5), N/A (13) | (2, 12) |
| *Streptococcus mitis* | 9.97, 12 | HPLC (9.97), N/A | (2, 72, 73) |
| *Sulfolobus* | 7 | Periodate oxidation | (62) |
| *Synechocystis* sp. PCC6803 | 7.5-10.4, 9.6 +/- 0.4 | High-Performance Anion-Exchange Chromatography | (74, 75) |
| *Thermococcus* | 7 | Periodate oxidation | (62) |
| *Thermoproteus* | 7 | Periodate oxidation | (62) |
| *Thermus thermophilus* | 7 | Anion-Exchange Chromatography with a pulse amperometric detector | (76) |

| Eukaryotes | gc | Methods | Ref |
| --- | --- | --- | --- |
| Bass liver | 14 | Periodate oxidation | (77, 78)) |
| Bullhead liver | 12 | Periodate oxidation | (77) |
| Cat liver | 13-15 | Periodate oxidation,<br>Isoamylase | (79) |
| Dog liver | 12 | Periodate oxidation | (77) |
| Haddock liver | 12 | Methylation | (78) |
| Horse liver | 11 | Periodate oxidation | (77) |
| Horse muscle | 11-12 | Methylation | (80) |
| Human muscle | 11-12 | Periodate oxidation,<br>Isoamylase | (79) |
| Northern pike liver | 12-13 | Periodate oxidation | (78) |
| Rabbit muscle | 11-13 | Periodate oxidation,<br>Methylation, Isoamylase | (79, 80) |

## Updated Evidence for Glycogen as a Durable Energy Reserve

Since the proposal of short *g_c_* glycogen as a durable energy reserve, more experimental evidence has come up to support the hypothesis indirectly (Table 1). A study by Grundel et al. noted that, compared with other energy reserves such as polyhydroxybutyrate (PHB), only glycogen plays the decisive role in metabolic adaptation and maintenance of Synechocystis viability during malnutrition conditions (7). In addition, the study also suggested that short *g_c_* glycogen has a durable energy reserve role in Cyanobacteria, which is essential for bacterial survival in harsh conditions (7). In the context of a study of Agrobacterium tumefaciens, Bains simply stated that short *g_c_* glycogen could be utilized slowly due to steric inhibition and promote bacterial survival under starvation (8). Finally, in a targeted metagenomic analysis of glucan-branching enzyme genes in fecal microbiota, Lee et al. emphasized that more work should be done on the effects of glycogen average chain length in order to better understand the functions of glycogen-degrading enzymes, i.e., how *g_c_* links with degradation rate (9).

Wang et al. was the first, and currently the only, group that tried to manipulate glycogen *g_c_* through altering in situ the N-terminus of glycogen branching enzyme (GBE) (6). Although it was proven to be feasible, this experiment brought in multiple confounding factors and could not reach a clear conclusion about how glycogen *g_c_* influences *E. coli* durability. Recently, Jo et al. identified that Vibrio vulnificus is able to accumulate very short *g_c_* glycogen and inferred that this feature may be associated with *V. vulnificus*’s unique life cycle in the marine environment (10). In addition, Han et al. used the hypothesis to explain the relationship between glycogen structure and function in their study (11). During the study of highly branched glycogen in red microalga *Galdieria sulphuraria*, Martinez-Garcia et al. initially presented the two theories that short *g_c_* glycogen could be an easily accessible energy supply or a very durable energy reserve without giving a preference (12). In the follow-up study, they confirmed that short *g_c_* glycogen is a slowly digestible and resistant glucose polymer (13), which provided a strong support for glycogen as a durable energy reserve. In a recent editorial, Klotz and Forchhammer also viewed positively the hypothesis based on the current knowledge (14). Other than these studies, most of the rest listed only cited the hypothesis to support the idea that bacteria with glycogen metabolic pathways are able to face diverse environments and have a flexible lifestyle, which is not relevant to our topic from this perspective (15, 16).

Although there have been no bacterial glycogen structure studies to date, progress has been achieved in higher organisms such as human and animals, according to which, glycogen could have different degradation patterns depending on structural differences. Recent analysis via size exclusion chromatography (SEC) and fluorophore-assisted carbohydrate electrophoresis (FACE) showed that rosette-shaped α- particles in liver glycogen are fragile and easily broken down into smaller β- particles, which degrade much faster than a-particles and may contribute to uncontrolled hyperglycemia (17, 18). However, the explanation about why α- particles are more resistant to degradation than β- particles is simply based on surface area to volume ratio and is debatable (19). In fact, if phosphorylase expression is up-regulated to sufficient concentration in the reaction, large particles should be degraded faster than smaller ones. Moreover, C^13^-labeled glucose study showed that brain glycogen synthesis in astrocytes is around 25 times slower than liver glycogen (20). In addition, abnormal glycogen structure has also been observed in different types of glycogen storage disease (GSD) cases, which generates great impacts on glycogen degradation and normally has adverse effects on human health (21).

In sum, glycogen structure is heterogeneous and not consistent, even in the same domain of life or in the same species, including human and animals. Recent research has been warming to the idea that short *g_c_* glycogen could have a potential role as a durable energy reserve. However, direct experimental evidence is needed in order to form a much clearer picture about the process in bacteria.

## Theoretical Modeling of Glycogen Structure

To further investigate our hypothesis, we have created theoretical models to describe the structure of glycogen and thus explore the important roles of *g_c_* in glycogen. Using the classical Whelan model (22), together with conditions derived from physical constraints and experimental data that we describe here, a set of glycogen theoretical structures were generated. Through comparative analysis, it was suggested that short *g_c_* glycogen has smaller size, higher percentage of α-1, 6-linkages, higher density in the outmost layer, and fewer phosphorylase available glucosyl residues, while the number of the non-reducing ends are the same. Thus, short *g_c_* glycogen is very likely hard to degrade. Specifically, the following eight mathematical formulas describe a glycogen particle in Whelan’s model (1). Glycogen particles include A- and B-chains and are divided into tiers (t). A-chains are those in the outmost tier without any branches while B-chains are distributed in interior with branches (degree of branching r). All chains have the same length *g_c_*, i.e., average chain length. Based on this simplified model, a set of mathematical expressions has been constructed to describe glycogen structure as follows(1).

1. Total number of chains in a glycogen particle CT

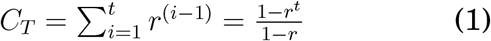

* r is the degree of branching, that is, the average number of branches in a B chain.
2. Number of chains in a given tier Cti

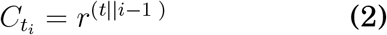

* ti is the i-th tier
3. The number of A-chains CA

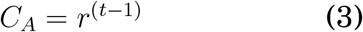
4. The number of glucose available to phosphorylase in a single A chain GPC

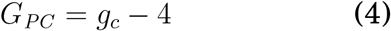 * The reasoning behind this equation is that phosphorylase can only break α-1, 4- glycosidic bonds one at a time from the non-reducing end of a chain until 4 glucosyl residues away from α-1, 6- glycosidic bonds (branching points)
5. The number of glucose available to phosphorylase in a glycogen particle GPT

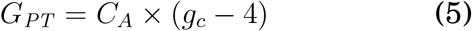
6. Total glucose in a glycogen particle GT

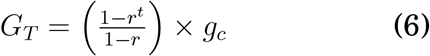
7. Radius of a glycogen particle Rs

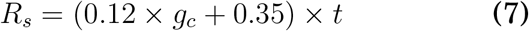
8. Volume of a glycogen particle Vs

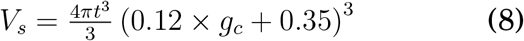

According to glycogen mathematical model (formulas listed above), theoretical spatial limitations of glycogen structure (physical constrains), and experimental data sourced from bacterial glycogen studies, we propose four constraints for bacterial glycogen structure, which are listed below.

1. The sum of cross-section area of glucose should not be greater than the surface area of glycogen sphere at the outmost layer.

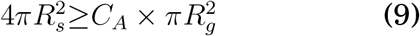
1. Glycogen tier volume should be greater than the total volume of glucosyl residues.

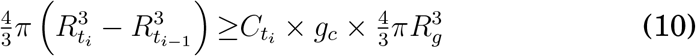
1. Molecular weight is between 107 to 108 Dalton, that is, the number of glucosyl residues should be in the range of 55506 and 555062 in a single glycogen molecule (23).

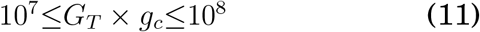
1. Glycogen diameter ranges from 20 to 50 nm (24).

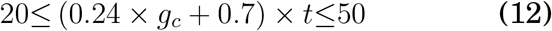

Through numerical calculation based on the four newly developed constraint conditions via R package, at given *g_c_* and r, we get the maximal tier number t in correspondence. For a complete list of maximal tier numbers, please see Table S1 in the Supplementary Data file. Specifically, for *g_c_* ranging from 7 to 12, the maximal tier is only 13. If *g_c_* is larger than 13, 2 or more maximal tiers could be achieved, depending on branching degrees. In order to understand how glycogen *g_c_* influences glycogen structure, we chose 3 representative *g_c_* in our study (shown with a * in Table S1), which are 7, 10, and 13 with r = 2 and t = 13. Glycogen with r = 2, t = 13, *g_c_* = 7, is the structure that is most likely present in bacteria, while r = 2, t = 13, *g_c_* = 13 is a typical glycogen structure identified in eukaryotes. By comparing these two structures, we can get a hint about how glycogen *g_c_* can influence its degradation. Thus, a theoretical three-dimensional model is constructed for each *g_c_* (Figure 1) and the corresponding parameters are presented for each model (Table S2).

**Figure 1:**
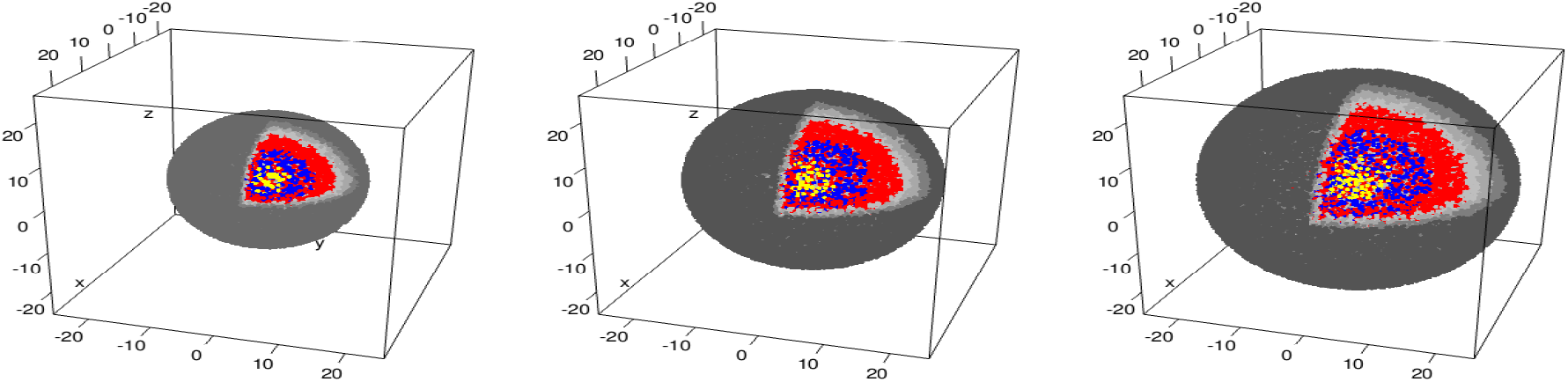
Three-dimensional illustration of bacterial glycogen structure at different gc values with given degree of branching r and maximal tier number *t*. Physical properties varies and the most apparent difference among them is the sphere diameter (*D*_*gc*=7_ = 28.56 nm, *D*_*gc*=10_ = 40.3 nm, *D*_*gc*=13_ = 49.66 nm). Although it is predicted that the ratio of surface (*S_s_*) and volume (*V_s_*) determines glycogen degradation rate, the conclusion does not consider glycogen interior structure. Thus, it would be interesting to see how degrading rate changes by combing the two factors to simulate glycogen degradation. Tier 1-3, shown in yellow dots; Tier 4-6 shown in blue dots; Tier 7-9 shown in red dots; Tier 10-13 shown as a gradient from light to dark grey.

Based on the structure modelling, glycogen with parameters r = 2, t = 13, *g_c_* = 7 has maximal diameter of D*g_c_* = 7 = 28.56 nm while glycogen with parameters r = 2, t = 13, *g_c_* = 13 has maximal diameter of D*g_c_* = 13 = 49.66 nm. Thus, short *g_c_* leads to small particle size. Further analysis shows that small glycogen particles (*g_c_* = 7) have a smaller number of glucose molecules available to glycogen phosphorylase at the outmost sphere than glycogen particles with *g_c_* = 13. Previous study confirmed that glycogen phosphorylase has a high affinity to glycogen if *g_c_* is longer (2). Recent investigation into acid hydrolysis of liver glycogen showed large α- particles degrade at faster rate than small β- particles (25), which may be due to more available glucosyl residues at the outermost shell. Thus, larger glycogen particles could be degraded more easily due to long *g_c_*. In addition, glycogen particle (*g_c_* = 7) is much denser than glycogen particle (*g_c_* = 13) at each tier, based on the calculation of *ρ* = *G_ti_*/(*Vs_ti_* − *Vs*_*ti*−1_). Higher density may exert spatial hindrance for the glycogen debranching enzyme to function (26). Because branching degree r is pre-set, no difference in terms of the number of α-1, 6- branching points is identified in the models. In sum, based on phosphorylase-available glucose and density in a glycogen particle, we can infer that short *g_c_* glycogen is harder to degrade.

## Glycogen Metabolism and its Relationships with Bacterial Environmental Survival

Experimental studies have provided considerable evidence to support glycogen’s contribution to bacterial survival in the environment under a variety of stresses, such as temperature fluctuation, nutrient deprivation, and osmolarity instability (14, 27–30). A variety of genes have been identified to be related with glycogen accumulation and bacterial environmental survival based on Keio collection and ASKA library analyses, such as csrA, rpoS, spoT, phoP, and phoQ, etc. (31, 32). A comprehensive review on bacterial glycogen can be found at Wilson et al. in 2010 (23), which revealed interconnections between glycogen metabolism and cellular processes. The most recent review about glycogen given by Park, however, focused on glycogen metabolism related enzymes (24). No much bacterial durability was mentioned. Thus, a need for an updated review about glycogen and environmental persistence in bacteria is urgently need. Here, we give an update about recent progress in terms of how glycogen metabolism may contribute to bacterial environmental survival, by focusing on the two compounds, maltodextrin and trehalose (Figure 2). Enzymatic reactions, rather than regulatory mechanisms, are concerned here.

**Figure 2:**
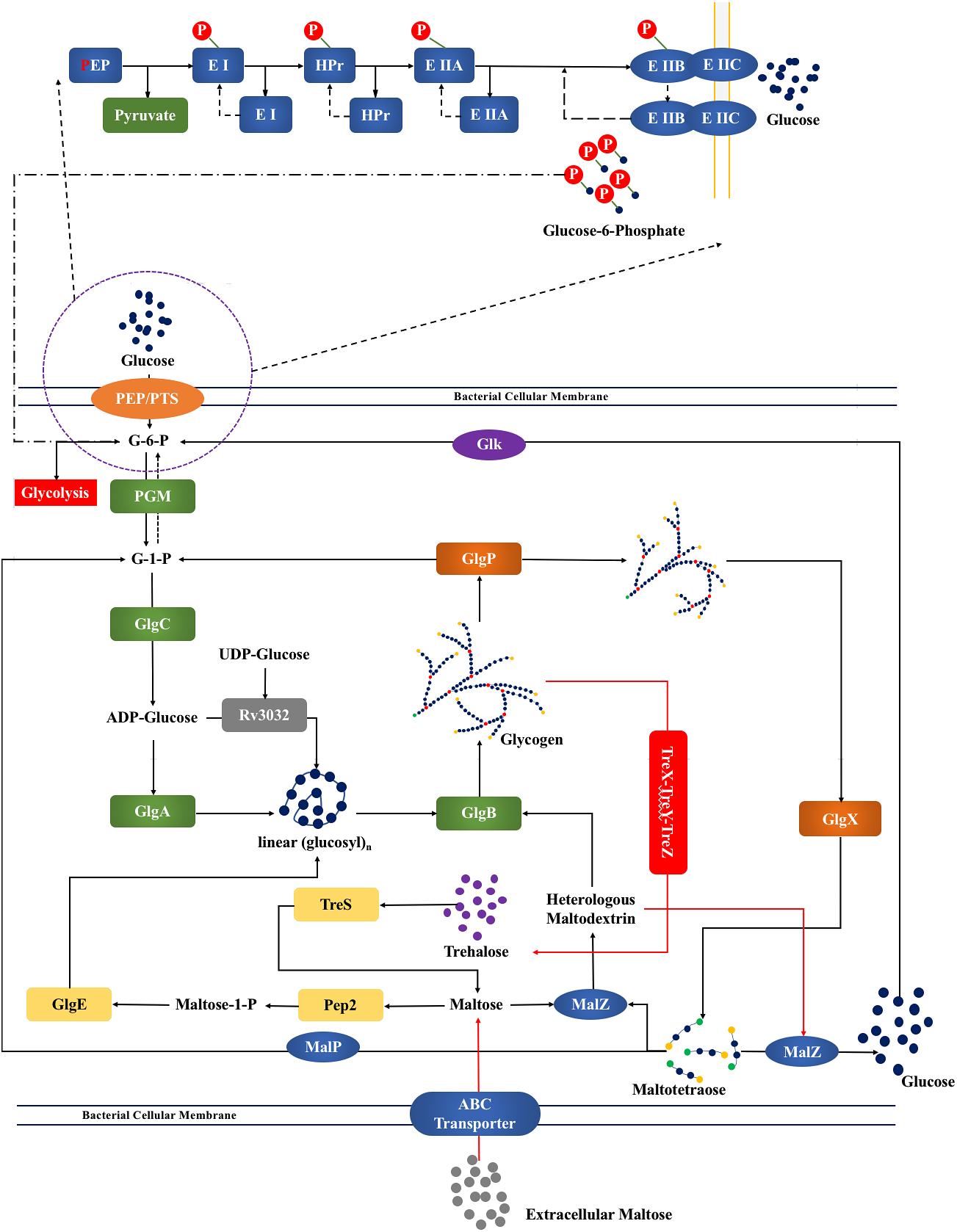
Bacterial glycogen metabolism with integrated maltose and trehalose pathways. Extracellular free glucose was phosphorylated into G-6-P through PEP:PTS system during transporting into cytoplasm. G-6-P was then transformed into G-1-P by PGM. GlgC provided ADPG as building blocks for GlgA. GlgB then branched the linear oligosaccharide into highly branched structure. Glycogen was degraded first by GlgP from outmost sphere to generate G-1-P and then truncated by GlgX to generate maltotetraose. The six enzymes work concurrently to balance glycogen metabolism and maintain its structure dynamically. Newly discovered associations of trehalose (TreS, Pep2, GlgE, GlgB) and maltose (MalQ, MalZ, MalP) pathways with glycogen metabolism are also illustrated. In addition, the classical TreX(GlgX)-TreY-TreZ pathway for trehalose synthesis from glycogen is also present, while another GalU-OtsA-OtsB pathway has been omitted (29). Details are in the main text. Rv3032 is another newly identified enzyme responsible for possible intracellular and capsular glycogen formation in *Mycobacterium tuberculosis*, exclusively. Glycogen, trehalose, and maltose are interconnected, forming a dynamic network for bacterial viability in the face of unfavourable conditions.

Maltodextrin metabolism was previously linked with bacterial osmoregulation, i.e,, sensitivity of endogenous induction to hyperosmolarity (30), which interacts with glycogen metabolism. For osmoregulation, glycogen-generated maltotetraose is dynamically metabolised by MalP (maltodextrin phosphorylase) and MalZ (maltodextrin glucosidase), while MalQ is responsible for maltose recycling to maltodextrins (30). Park et al. looked into the metabolism and proposed a molecular mechanism for the relationship between glycogen and maltose metabolism (33). Basically, maltotetraose generated via glycogen branching enzyme truncation will be further processed by MalQ for glycogen recycling by forming maltodextrin or by MalZ for glycolysis of glucose with the assistance of glucokinase (Glk) (24). Otherwise, glucose-1-phosphate is formed through MalP as a building block of glycogen synthesis or for glycolysis (24). Thus, extracellular maltose or maltodextrin derived from glycogen degradation are tightly linked with glycogen metabolism and have the potential as an essential player for bacterial environmental persistence. In addition, maltose may participate in the formation of capsular α-glucan through (TreS)-Pep2-GlgE-GlgB pathway, which plays essential roles for environmental persistence and antibiotic resistance of Mycobacterium tuberculosis (34, 35).

Trehalose plays essential roles in bacterial adaptation to temperature fluctuation, hyperosmolarity, and desiccation resistance (28, 36). It was well-established that trehalose can be generated from glycogen through the TreX(GlgX)-TreY-TreZ pathway (37), or from the GalU-OtsA-OtsB pathway (38). Recently, an unexpected and widespread connection between trehalose and glycogen (TreS-Pep2-GlgE-GlgB) has been identified in bacteria (29), which provides a complementary way of cycling these two metabolites. Dalmasso et al. confirmed that both glycogen and trehalose are accumulated under cold conditions (4°C) in *Propionibacterium freudenreichii* species (28). Although it was suggested that the two compounds provide a molecular basis of the long-term survival and activity during prolonged incubation at low temperatures, no specific mechanisms were provided. Reina-Bueno systematically examined the roles of trehalose in *Rhizobium etli* in terms of abiotic stress resistance (36). Enhanced gene expression (otsA and otsB) and trehalose content were observed. However, linkages between glycogen and trehalose metabolism was not mentioned. In addition, it is also noteworthy that TreT, a trehalose synthase, sporadically identified in a small set of bacteria (*Rubrobacter xylanophilus*) and archaea (*Pyrococcus furiosus*), catalyses the reversible formation of trehalose based on ADP-glucose and glucose (23). Thus, TreT in trehalose metabolism might contribute to glycogen biosynthesis, although further experimental validation is necessary.

Glycogen plays essential roles in bacterial energy metabolism and is widely connected with a variety of metabolic pathways that are tightly associated with bacterial persistence in the face of environmental stresses such as starvation, desiccation, temperature fluctuation, and/or hyperosmolarity. Maltodextrin and trehalose pathways are examples. However, how glycogen itself and its structure exert impacts on bacterial persistent phenotypes through the connection with other compounds require further exploration and might shed light on new mechanisms for understanding bacterial persistence strategies.

## Conclusions

It has been established that glycogen is an important energy reserve in bacteria (23). In this perspective, we have provided an updated view of recent progress in the study of glycogen as durable energy reserve in bacteria since the DESM hypothesis was proposed (2). Recent relevant literature has reinforced the feasibility of the hypothesis. A preliminary glycogen structure modeling also provided a theoretical support for the hypothesis. Ideally, the structure constraint will force bacteria to utilize glycogen in a controlled manner, i.e., slow release of glucosyl residues, which may contribute the elongated survival of bacteria in nutrient-deprived or other life-challenging conditions. Although the network of glycogen metabolism has been investigated from genome (Keio collection) (31), transcriptome (ASKA library) (32) and systematic levels (Petri net model) (39), knowledge about the associations between glycogen metabolism and bacterial environmental persistence is still limited. An updated view about how these genes interact with bacterial environment durability and how glycogen structure can impact on bacterial persistence would greatly improve our knowledge of glycogen in bacterial physiology. Taken together, priority at current stage should be given to experimental investigation of how *g_c_* influences glycogen degradation. Then, the next questions are whether slow degradation of glycogen could prolong bacterial viability and what are the specific mechanisms. By answering these questions through well-designed experiments, we would get more insight into relationship between bacterial glycogen structure and its biological activities.

## Acknowledgements

We would like to thank the Startup Foundation for Excellent Researchers (2016) provided by Xuzhou Medical University under grant number D2016007, the Nature and Science Foundation for Colleges and Universities (2016) by Jiangsu Province under grant number 16KJB180028, Innovative and Entrepreneurial Talent Scheme of Jiangsu Province (2017), and Natural Science Foundation of Jiangsu Province (2018). Dr. Liang Wang is very grateful to Academician/Professor Robert G. Gilbert from University of Queensland and Yangzhou University for his constructive discussions.

