## Supplementary Data for "Bacterial Glycogen as a Durable Energy Reserve Contributing to Persistence: An Updated Bibliography and Mathematical Model"

**Table S1** Maximal tier number for a pair of given $g_c$ and $r$ under four constraint conditions.

| $g_c$ | $r$=2 | $r$=3 | $r$=4 | $r$=5 |
|---|---|---|---|---|
| **7***| 13 | - | - | - |
| **8** | 13 | - | - | - |
| **9** | 13 | - | - | - |
| **10*** | 13 | - | - | - |
| **11** | 14 | - | - | - |
| **12** | 13 | - | - | - |
| **13*** | 13 | 8 | - | - |
| **14** | 12 | 8 | - | - |
| **15** | 11 | 8 | - | - |
| **16** | 11 | 8 | - | - |
| **17** | 10 | 8 | 6 | - |
| **18** | 9 | 9 | 7 | - |
| **19** | 9 | 9 | 7 | - |
| **20** | 9 | 9 | 7 | - |
| **21** | 8 | 8 | 7 | 6 |

Note: Although the space limitation between branching points is set to be 4 glucosyl residues, some doubly branched pieces are found, indicating that the space could be smaller and is variable [1]. As for $g_c$=7 and 8, the situation is clear that only 3 glucosyl residues are present between branching points so as to form 2 branches in an A chain. $g_c$ value marked with * will be used for glycogen modelling due to 1) $g_c$ difference and 2) same $r$ and maximal $t$.
